# Epithelial-mesenchymal plasticity induced by discontinuous exposure to TGFβ1 promotes tumour growth

**DOI:** 10.1101/2021.11.02.466914

**Authors:** Mafalda Santos, Marta Ferreira, Patrícia Oliveira, Nuno Mendes, Ana André, André F. Vieira, Joana B. Nunes, Joana Carvalho, Sara Rocha, Mafalda Azevedo, Daniel Ferreira, Inês Reis, João Vinagre, Joana Paredes, Alireza Heravi-Moussavi, Jorge Lima, Valdemar Máximo, Angela Burleigh, Calvin Roskelley, Maria de Fátima Carneiro, David Huntsman, Carla Oliveira

**Affiliations:** I3S, Instituto de Investigação e Inovação em Saúde - Rua Alfredo Allen, 208, 4200-135 Porto, Portugal; Ipatimup, Instituto de Patologia e Imunologia Molecular da Universidade do Porto- Rua Júlio Amaral de Carvalho, 45, Porto, Portugal; INEB, Instituto de Engenharia Biomédica - Rua Alfredo Allen, 208, 4200-135 Porto, Portugal; British Columbia Cancer Agency - 600 W 10th Avenue, Vancouver, BC V5Z 4E6, Canada; Department of Pathology and Oncology of the Medicine Faculty of the University of Porto- Alameda Prof. Hernâni Monteiro, 4200 - 319 Porto, Portugal

**Author notes:** Correspondence; Expression regulation in Cancer Group, Lab 208S4 - i3S, Instituto de Investigação e Inovação em Saúde, Universidade do Porto, Rua Alfredo Allen, 208, 4200-135 Porto, Portugal. Co-authors (Mafalda Santos, Marta Ferreira). **Simple Summary:** In this manuscript, we used a non-genetically manipulated EMT/MET cell line model to demonstrate that epithelial mesenchymal plasticity occurring in normal cells generates co-existing phenotypically and functionally divergent cell subpopulations which result in fast growing tumours *in vivo*.

**Keywords:** MET, Cellular Heterogeneity, Self-Renewal, EMT, tumorigenic potential

## Abstract

Transitions between epithelial and mesenchymal cellular states (EMT/MET) contribute to cancer progression. We hypothesize that EMT followed by MET promotes cell population heterogeneity favouring tumour growth. We developed an EMT model by on/off exposure of epithelial EpH4 cells (E-cells) to TGFβ1 that mimics phenotypic EMT (M-cells) and MET. We aimed at understanding whether phenotypic MET is accompanied by molecular and functional reversion back to epithelia, by using RNA sequencing, Immunofluorescence (IF), proliferation, wound healing, focus formation and mamosphere formation assays, as well as cell-xenografts in nude mice. Phenotypic reverted-epithelial cells (RE-cells), obtained after MET induction, presented pure epithelial morphology and proliferation rate resembling E-cells. However, RE transcriptomic profile and IF staining of epithelial and mesenchymal markers revealed a unique and heterogeneous mixture of cell-subpopulations, with high self-renewal ability fed by oxidative phosporylation. RE-cells heterogeneity is stably maintained for long periods after TGFβ1 removal, both *in vitro* and in large derived tumours in nude mice. Overall, we show that phenotypic reverted-epithelial cells (RE-cells) do not return to the molecular and functional epithelial state, present mesenchymal features related with aggressiveness and cellular heterogeneity that favour tumour growth *in vivo*. This work strengthens epithelial cells reprogramming and cellular heterogeneity fostered by inflammatory cues as a tumour-growth promoting factor *in vivo*.

## Introduction

Epithelial to mesenchymal transition (EMT) and the reverse process, mesenchymal to epithelial transition (MET), are biological mechanisms naturally occurring during embryogenesis and regeneration [1, 2]. Although contradictory at first, the role of EMT and MET in cancer progression and metastization has now been fully acknowledged [2–6]. While EMT enables epithelial cancer cells dissemination, bestowing cells with increased invasion, migration and stem-cell properties, MET facilitates the establishment of these cells at secondary sites [3, 7, 8]. EMT and MET were previously seen as strict transition states where cells acquired specific phenotypes and molecular signatures. However, this biological programme is very dynamic and cannot be accurately defined by limited sets of markers or phenotypic changes. Concomitant expression of epithelial and mesenchymal markers in cancer cells suggests occurrence of hybrid EMT states [1,8–10]. This cellular plasticity confers advantageous features to cancer cells conferring them with increased adaptability to microenvironment cues and resistance to several stressors [11]. Supporting this, Armstrong et al showed that >75% of circulating tumour cells (CTCs), isolated from patients with metastatic prostate and breast cancers, exhibited intermediate phenotype and stem-cell markers [12]. Moreover, Yu et al observed that CTCs from breast cancer patients, show a varying proportion of epithelial/mesenchymal markers associated with different breast cancer subtypes and treatment responses [13].

Many strategies have been described to induce EMT *in vitro*, such as artificially-induced overexpression of transcription factors, such as Snail and Twist1 [14–16] or treatment with growth factors/cytokines, such as TGFβ1, EGF and NGF [1,17,18]. These *in vitro*, as well as *in vivo* studies have strengthened the hypothesis that EMT followed by MET occurs at different levels of cancer progression. Hugo et al showed that primary tumours derived from breast cancer cells, exhibited evidences of EMT at the invasive front, while derived metastasis expressed high levels of E-cadherin, suggesting MET [8]. Tsai et al showed that after activation of the EMT-inducer Twist1, cancer cells disseminated into the blood circulation, but Twist1 was inactivated to induce MET, allowing disseminated cancer cells to metastasize [19]. In line with this, Ocaña et al demonstrated that loss of the EMT-inducer Prrx1, together with the acquisition of an epithelial phenotype and stem-cell properties, were required for cancer cells to form metastases *in vivo*, reinforcing MET as an important event for cancer colonization [20].

The role of EMT and MET is currently well established in tumour progression and several reports also correlate EMT-drivers with increased stemness and prevalence of tumour-initiating cells (TICs) [21,22]. Current EMT/MET models always imply either the study of mammary stem cells and cancer stem cells separately or they promote cell transformation through activation of an oncogene, such as KRAS [22]. The EMT driver TGFβ was able to trigger increased breast TICs in claudin-low breast cancer cell lines [23]. However, in a different study, it was suggested that TGFβ had an inhibitory role in breast TICs [24]. The molecular context in which these events occurred were not disclosed.

Herein, we hypothesize that EMT followed by MET promotes cell-population heterogeneity, and that this favours tumour growth. We characterized and explored an EMT/MET model and unveiled that MET generates population heterogeneity, which may drive tumour growth *in vivo*.

## Results

### Phenotypic EMT/MET *vs* molecular EMT/MET

To decipher the mechanisms underlying naturally-occurring MET, we established and characterized an *in vitro* EMT/MET model using the near-normal EpH4 mouse mammary epithelial cell line (E-cells) exposed to the EMT-inducer TGFβ1 [25]. This non-cancer cell line was selected to prevent cancer-related bias. Moreover, this model has an homogenous nature both in terms of brightfield morphology and epithelial/mesenchymal markers expression [25]. After 7-day of TGFβ1-treatment, E-cells acquired a fibroblastoid phenotype, resembling mesenchymal cells (M-cells, Fig 1a). TGFβ1 was then removed from the culture medium and after another 4 days, brightfield microscopy revealed widespread recovery of an epithelial phenotype (Reverted-Epithelial, RE-cells, Fig 1a, Supplementary Fig 1).

**Fig 1.**
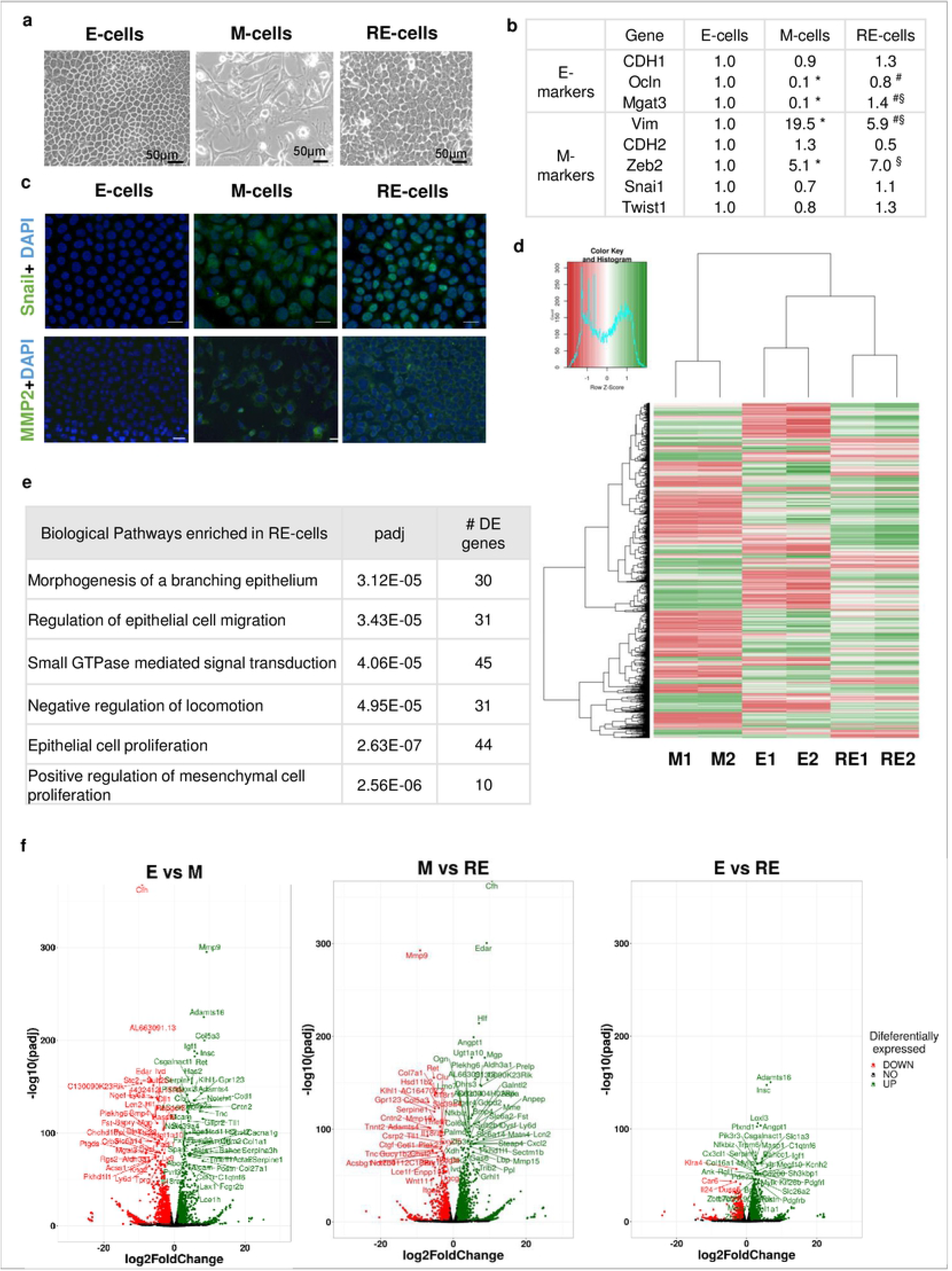
RE-cells underwent MET and display a transcriptomic signature that partially resembled E-cells. **(a)** Brightfield images of E-, M-, RE-cells. **(b)** RNA expression analysis of EMT-associated markers in E-, M-, RE-cells (qRT-PCR, * for *p*<5.00E-02 for E-*vs*. M-cells, § for *p*<5.00E-02 for E-*vs.* RE-cells, # for *p*<5.00E-02 for M-*vs.* RE-cells). **(c)** Immunofluorescence staining for Snail, MMP2 and DAPI as indicated. RE-cells show a strong staining for both Snail and MMP2, indicating that they retain M-features. **(d)** Heatmap of differentially-expressed genes assessed by RNAseq (0.6>fold-change>1.5, *p*<1.00E-02). Z-scored values from −1 to 1 (red to green, from low to high expression levels). (**e)** Significantly-enriched biological pathways derived from the 1288 differentially-expressed genes specific for RE-cells. (**f)** Volcano Plots highlighting the differentially expressed genes in all the comparisons among the different cell lines.

To confirm EMT-induction/reversion, E-, M- and RE-cells were characterized for expression of epithelial (*CDH1, Ocln, Mgat3*) and mesenchymal (*Vim, CDH2, Zeb2, Snai1, Twist1*) markers by qRT-PCR (Fig 1b). As we previously reported, we did not observe a significant alteration in *CDH1* expression, however the function of the corresponding protein E-cadherin was impaired, due to downregulation of Mgat3 expression in M-cells, which is responsible for GnT-III-mediated glycosylation [25]. Moreover, M-cells displayed significant downregulation of other epithelial markers (Ocln) and upregulation of mesenchymal markers (Vim, Zeb2). In RE-cells, the expression of Ocln, Mgat3, Vim returned to levels similar to those of E-cells. In contrast, Zeb2 expression remained elevated in RE-cells, while Snai1 and Twist1 exhibited no alterations across E-, M- and RE-cells (Fig 1b). Immunofluorescence staining of these E-, M- and RE-cells for Snail and MMP2 mesenchymal markers showed the classical EMT pattern, since they are not expressed in E-cells and are expressed in M-cells, but remained unchanged in RE-cells as compared to M-cells, suggesting that RE-cells did not fully revert to the epithelial state (Fig 1c). Overall, the phenotypic changes, gene expression and immunofluorescence results support that the current EMT model mimics phenotypic EMT (M-cells) and MET, but also suggests that phenotypic MET may not be accompanied by molecular and functional reversion back to epithelia.

### Phenotypic MET is not supported by complete molecular reversion back to epithelia

We next used whole transcriptome sequencing (RNAseq) to explore differences and similarities between E-, M- and RE-cells. A good correlation was observed between the expression pattern obtained through qRT-PCR and RNAseq for epithelial/mesenchymal markers (Supplementary Fig 2). This validation allowed the use of RNAseq data to assess the expression variation of other EMT-associated markers, that supported EMT and partial MET (Supplementary Fig 2). RNAseq data was also used to identify Differentially Expressed Genes (DEGs) across the transcriptomic landscapes of E- and M- and RE-cells, by comparing: 1) E- and M-cells; 2) M- and RE-cells and; 3) E- and RE-cells (fold-change>1.50 or <0.66, p<1.00E-02, Supplementary Fig 3). These DEGs, were then submitted to double hierarchical clustering analysis (Fig 1d). Although, RE-cells signature was overall more closely related to E- than to M-cells, this cell state presents its own transcriptomic landscape. Indeed 1288 genes expressed in RE-cells, significantly changed specifically in this cellular state, while remaining stable during EMT (both in E- and M-cells). Among the top-ranking biological pathways there were ‘morphogenesis of a branching epithelium’, ‘regulation of epithelial cell migration’, ‘small GTPase mediated signal transduction’, but also ‘negative regulation of locomotion’. Both ‘epithelial cell proliferation’ and ‘positive regulation of mesenchymal cell proliferation’ were also part of highly significant biological pathway in RE-cells (Fig 1e). These data further support that MET generated epithelial-looking cells, differ from their original epithelial counterpart at the molecular level, but also that they differ significantly from the M-state cells from which they arose (Fig 1d, e, f Supplementary Fig 4).

A deeper analysis of biological functions and pathways significantly-enriched across the experiment (Supplementary Fig 5), showed that “Cellular Growth and Proliferation”, “Migration”, “Metabolism”, “Stemness” and, “Cancer” were affected in these transitions. These observations further supported the molecular differences between E-, M- and RE-cells, highlighting that even though our i*n vitro* model was developed using a near-normal cell line, a significant association with aggressiveness and cancer-related features was detected upon EMT/MET induction.

### Phenotypic MET generates E-like, M-like and novel cellular subpopulations

Given that RE-cells seem to be transcriptionally heterogeneous and have a set of quite specific molecular features, we next assessed *in situ*, the immuno-expression of the epithelial marker E-cadherin and the mesenchymal marker Fibronectin in E-, M- and RE-cells (Fig 2a). E-cells displayed homogenous E-cadherin membrane staining and lacked Fibronectin, while M-cells showed an irregular staining of membranous E-cadherin (described in [25]) and high expression of extracellular Fibronectin (Fig 2a). RE-cells revealed a far more complex expression pattern of these two markers that evidenced the existence of four distinct sub-populations (Ecad+/Fn+; Ecad+/Fn-; Ecad-/Fn+; Ecad-/Fn-), some of them previously absent from E- and M-cells (Fig 2a). The most striking of all RE-cell populations, were those lacking E-cadherin expression (Ecad-/Fn+; Ecad-/Fn-), which appeared exclusively in RE- cells, a molecular change that is generally associated with EMT. To better assess the extent of RE-cell’s phenotypic heterogeneity, full slides of stained RE-cells were scanned. Of notice, no field was homogeneous for any of the four RE-subpopulations previously described, reinforcing their spatial co-existence (Supplementary Fig 6). To understand whether this heterogeneity was temporary and part of the reversal process back to the E-state, RE-cells were cultured for longer periods without TGFβ1. After 10 days, the same four sub-populations were still observed in RE-cell cultures (Fig 2b). We could confirm this was not a specificity of the EpH4 on/off model, as the same transdifferentiation protocol applied to the human immortalized normal breast epithelial cell line MCF10A cells, returned similar results. Brightfield images of MCF-10A E-, M- and RE-cells show that RE-cells are mixture of E-like and M-like cells (Fig 2c, upper panel). We could not optimize the Fibronectin staining in this cell line, so we chose Vimentin as a mesenchymal marker. Unlike EpH4 M-cells, MCF10A M-cells completely loose E-Cadherin expression (Fig 2c, bottom panel). On the other hand, MCF10-A cells behave similarly to EpH4 cells after TGFβ removal from the media, regarding heterogeneous E-Cadherin and Vimentin staining across the culture, confirming that upon TGFβ on/off exposure some cell populations do not fully revert to E-state.

**Fig 2.**
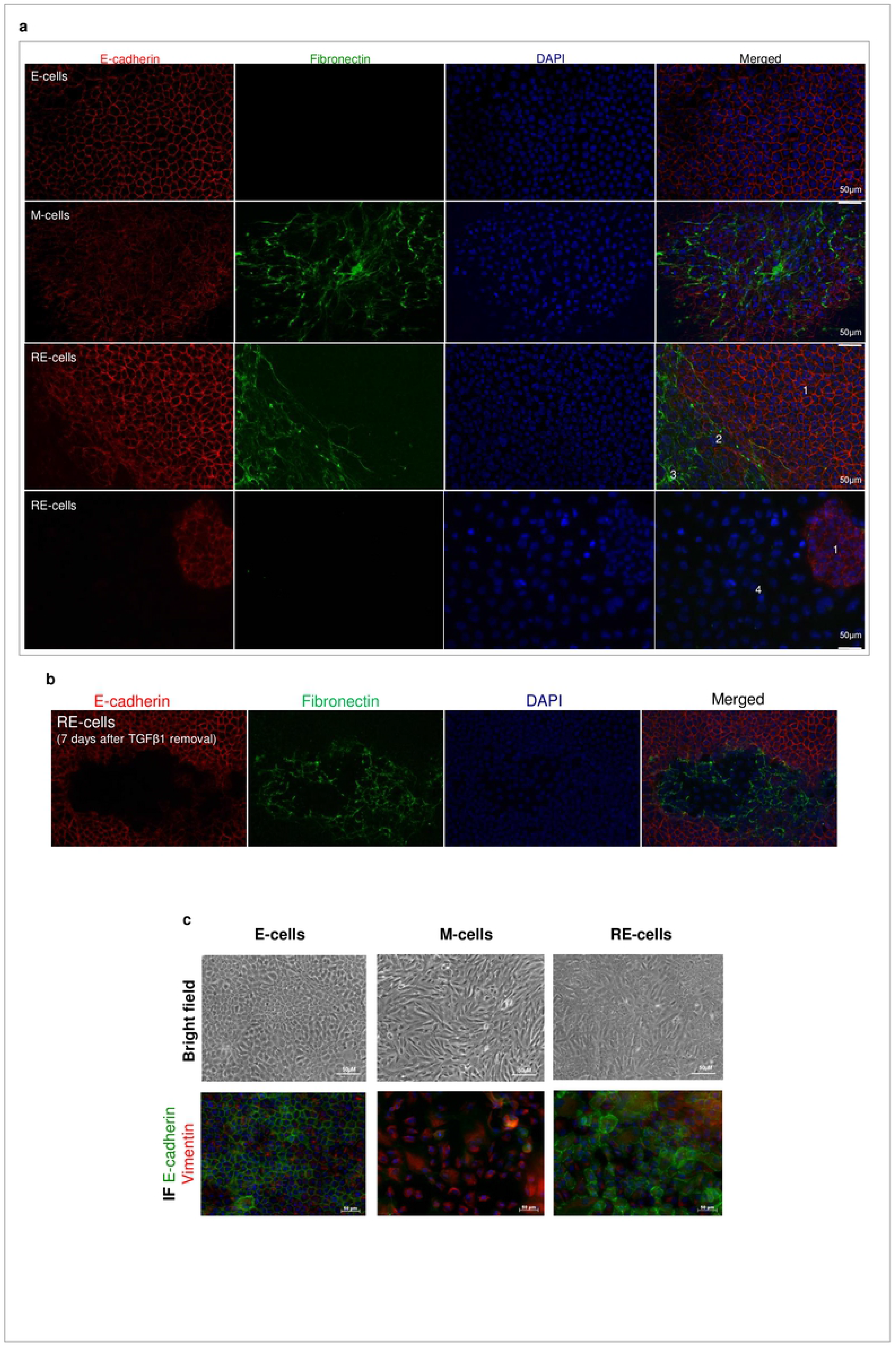
RE-cells are a mixture of different cellular subpopulations. **(a)** Immunofluorescence staining for E-cadherin (red) and Fibronectin (green) and DAPI (blue) of E-, M- and RE-cells. Immunofluorescence images highlighting RE-cells heterogeneity, according to the graph displaying E-Cadherin and Fibronectin intensities across slides **(b)** RE-cells retain their E-cadherin/Fibronectin staining heterogeneity even if reversion time is longer than 4 days. Immunofluorescence images for RE-cells grown for 7 day after TGFβ1 removal from the culture medium. **(c) Upper panel:** Brightfield images of MCF10A E-, M- and RE-cells. **Bottom panel:** Immunofluorescence staining for E-cadherin (Green) and Vimentinn (red) of MCF10A E-, M- and RE-cells.

### Cellular heterogeneity, generated after phenotypic MET in vitro, creates functional heterogeneity

Given that RE-cells represent an heterogeneous cellular population which is stable for several days in culture, we explored RE-cells functional behaviour in comparison to E- and M-cells. Moreover, we wanted to test the hints of aggressiveness observed in the transcriptomics analysis, eventually triggered by these transitions. For that, we analysed cell proliferation, cell behaviour when growing into a wound, and growth pattern in a focus formation assay, as several related biological functions were found enriched in RE-cells in the transcriptomics analysis (Fig 3a; Supplementary Fig 5). BrdU incorporation revealed that E- and RE-cells displayed a higher proliferation rate than M-cells, but only RE-cells were statistically different from M-cells (49%, 52% for E- and RE- *vs*. 34% for M-cells, *p*<0.05, Fig 3b). In fact, when comparing RE-cells with either M- or E-cells for proliferation-related DEGs, M-cells present a larger number of downregulated proliferation-associated genes than E-cells when both are compared with RE-cells, likely explaining why only M-cells differ from RE-cells in *the vitro* experiment (Fig 3b; Supplementary Fig 5).

**Fig 3.**
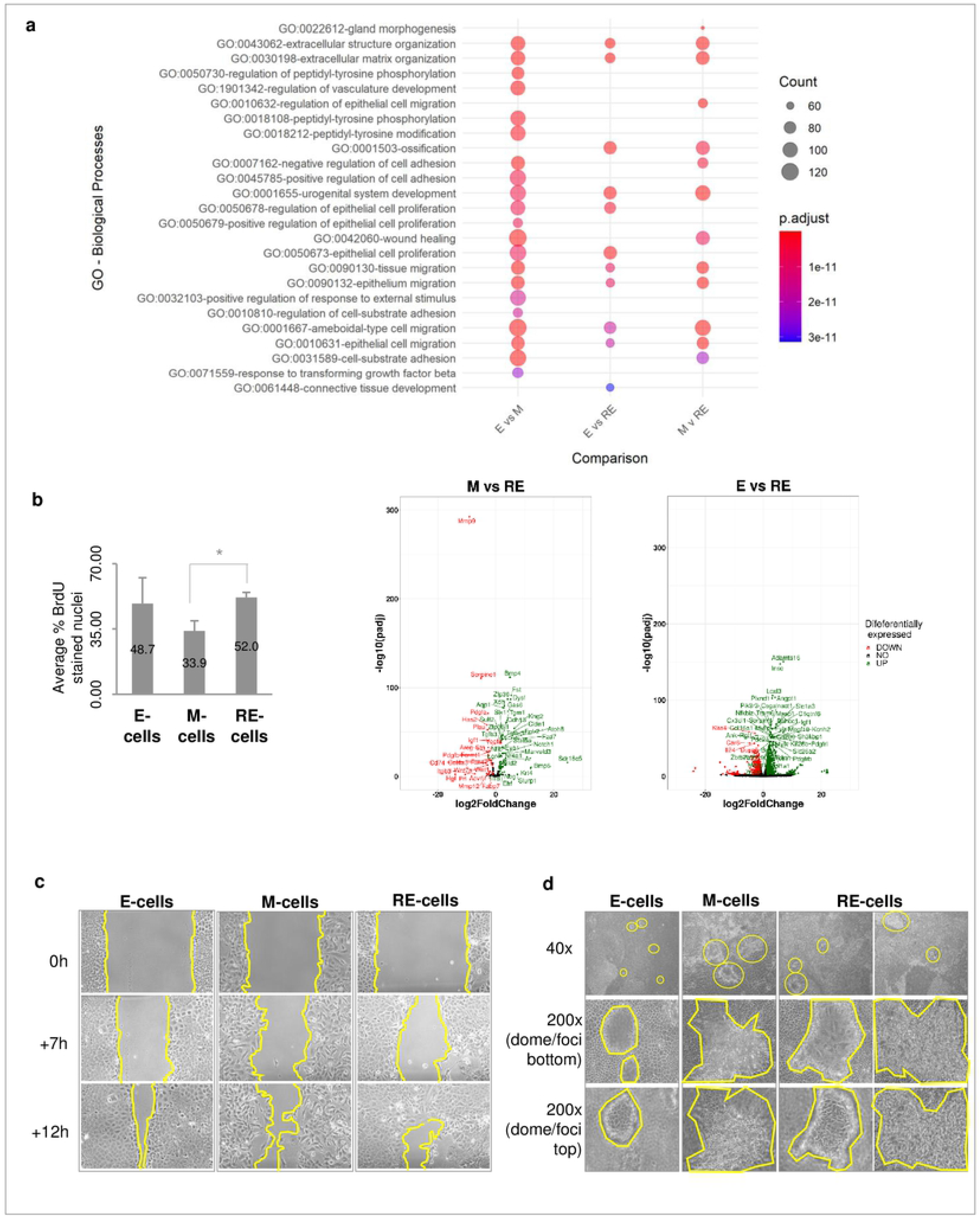
RE-cells exhibit high proliferation rate, heterogeneous migration pattern and focus formation assay. **(a)** Biological processes deregulated across comparisons (padj<0.05) displaying that RE-cells have a unique biological identity different from both E- and M-cells **(b)** Cell proliferation analysis, in terms of percentage of BrdU stained nuclei (per total number of DAPI-stained nuclei, n=3, \**p*<0.05 for M*vs*RE comparison).Volcano Plots showing upregulation of proliferation-related genes in RE-cells when compared to M- and E-cells.. **(c)** RE-cells exhibit mixed cell migration patterns, resembling both E and M-cells. Wound-healing brightfield images of E-, M-, RE-cells taken at different timepoints (0, 7, 12 hours, 100x). **(d)** Brightfield images of E-, M- and RE-cells grown for 21 days in plastic and normal culture medium. Top panel, general view of the 21-day-cultured E-, M- and RE-cells (40x). Middle panel, bottom layer of non-transformed cells surrounding dome-like structures or foci (200x). Bottom panel, top layer of dome-like structures or foci obtained after 21 days of culture of E-, M- and RE-cells (200x).

The wound-healing assays photographed and analysed at several timepoints showed that M-cells in the wound were mainly isolated, while a sheet of seemingly epithelial cells covered the wound area in E-cells (Fig 3c). In RE-cells, we observed both isolated cells, resembling those seen in M-cells, as well as areas with high cellular density, resembling the epithelial sheets seen in E-cells. In summary, RE-cells proliferated similarly to E-cells and faster than M-cells, while displaying both isolated cells and epithelial sheets covering the wound, reminiscent from both E- and M-cells. (Fig 3c).

To understand whether RE-cells acquired aggressive cancer-like features (Supplementary Fig 5), we next ran a focus formation assay. For this, E-, M- and RE-cells were cultured for 21-days and morphological differences were evaluated by brightfield microscopy (Fig 3d). E-cells generated few, small and spherical structures with defined edges, which, according to Gordon et al, could be considered dome-like structures resembling non-malignant mammary glands [26]. M-cells displayed a high number of foci, with large and irregular edges, but not dome-like structures, which is an indicator of increased aggressiveness. RE-cells displayed both E-like domes and M-like foci. Together with the previous data, this supports RE-cells as an entity with unique and heterogeneous phenotypes, retaining both E-like and M-like features in the population. These results supported the hypothesis that MET may confer a more aggressive phenotype to otherwise immortalized normal cells.

### Cellular heterogeneity, generated after phenotypic MET in vitro, is maintained in tumours growing in vivo

Our RNAseq data suggests that RE-cells are enriched in deregulated cancer-related pathways when compared to their E- and M-counterparts (Supplementary Fig 5). Therefore, we performed an *in vivo* pilot study, where cells that underwent EMT and MET were inoculated in the mammary fat-pad of athymic nude mice. Of notice, EpH4 cells have been described as non-tumourigenic [27]. In a pilot study, one out of two (1/2) mice inoculated with E-cells developed a tumour (6 mm3 at day 145) and 1/2 mice inoculated with M-cells developed another tumour (21 mm3 at day 132) (Supplementary Fig 7a). Both mice inoculated with RE-cells developed larger tumours (121 and 63 mm3 at day 145, Supplementary Fig 4a). This pilot study prompted us to assess the tumourigenicity of E-, M- or RE-cells in a larger group of mice (n=5, Fig 4a). In this second study, all cell types inoculated formed tumours, although with significantly distinct volumes. By the end of the experiment, E-tumours were significantly smaller in size (<30 mm^3^) than M-tumours (32-343 mm^3^) and RE-tumours (5-304 mm^3^, Fig 4b, c) (*p*<0.05). M- and RE-tumours were similar in size. All tumours were classified as malignant sarcomatoid carcinomas upon histopathological evaluation (Fig 4d). All tumours presented evidences of hyalinization, while only M- and RE-tumours displayed necrotic areas. M-tumours revealed signs of inflammation and local epidermis invasion while RE-tumours displayed increased cellular density (Fig 4d). The presence of mitotic nuclei and the different tumour volumes observed, led us to assess proliferation in E-, M- and RE-tumours by Ki67 immunostaining, however no significant differences were observed (Fig 4e, f, Supplementary Fig 7b-h).

**Fig 4.**
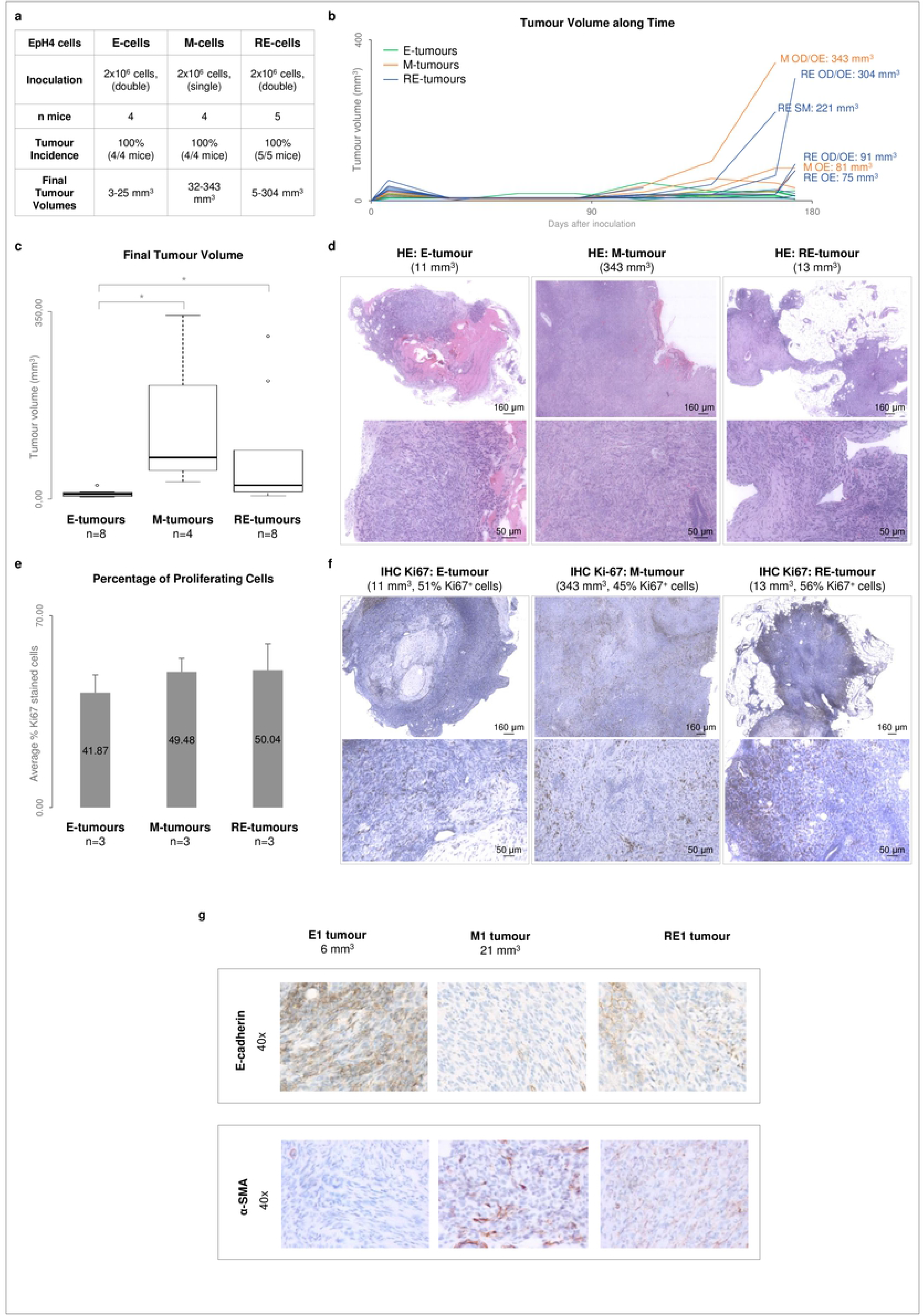
M and RE-cells display higher in vivo tumorigenicity that E-cells, without retaining the original *in vitro* RNA profile. **(a)** Summary of the *in vivo* tumorigenicity assay performed with E-, M- and RE-cells. **(b)** Growth curves representing the tumour volumes along time (E-tumours in green, M-tumours in orange and RE-tumours in blue). **(c)** Final tumour volumes obtained for each cell type (* for *p*<5.00E-02). **(d)** Representative images of hematoxylin and eosin staining of E-, M-, and RE-tumours. Top and bottom images with different magnification. **(e)** Average percentage of cells positive for Ki67 staining in 3 E-, 3 M- and 3 RE-tumours. **(f)** Representative images of immunohistochemistry staining for Ki67. Top and bottom images with different magnification**. (g)** E-cadherin and α-SMA immunohistochemistry staining of representative E-, M- and RE-tumours.

The molecular differences of E-, M- and RE-tumours were assessed through immunohistochemistry against E-cadherin and α-SMA. E-tumours expressed E-Cadherin but not α-SMA, whereas M-tumours lost E-cadherin expression and presented α-SMA staining. RE-tumours, however, expressed both markers, suggesting that the cellular heterogeneity created *in vitro* after MET, could be maintained after long periods *in vivo*.

### RE-cells mimic M-cells in self-renewal capacity in vitro and fast growth of tumour transplants in vivo

Our *in vitro* RNAseq data also revealed that not only M-but also RE-cells were enriched in stem-cell related pathways, as compared to E-cells (Suplementary Fig 5, Fig 5a), in agreement with literature showing that EMT generates cells with increased stemness [18]. Therefore, we explored whether M- and RE-cells displayed increased self-renewal capacity *in vitro*, using a first-passage mammosphere assay to evaluate whether E-, M- and RE-cells were able to grow in anchorage-independent conditions. Both M- and RE-cells displayed an increased ability to form first-passage mammospheres in comparison to E-cells (Fig 5b). Notably, RE-cells were able to form first-passage mammospheres with the same efficiency as M-cells (Fig 5b). In parallel, we tested the self-renewal ability of E-, M- and RE-tumours by syngeneic transplantation of small tumour fragments [28]. The histology of transplanted tumours mimicked that of the original tumours, however the growth rate of transplanted tumours was higher than the original counterparts (180 in the original experiment *vs* 49-84 days after transplantation) (Fig 5c). In particular, both M- and RE-transplanted tumours displayed a growth rate higher than the E-transplanted tumour, likely due to increased self-renewal ability [29]. In particular, two re-implanted fragments of an RE-tumour started their exponential growth just 15 days post-inoculation, when the original tumour took 110 days to start growing (Fig 5d, left panel). Immuno-staining of these tumours with E-cadherin α-SMA, revealed similar expression patters to that of the original tumours, further supporting that cellular heterogeneity is stable (Fig 5d, right panel).

**Fig 5.**
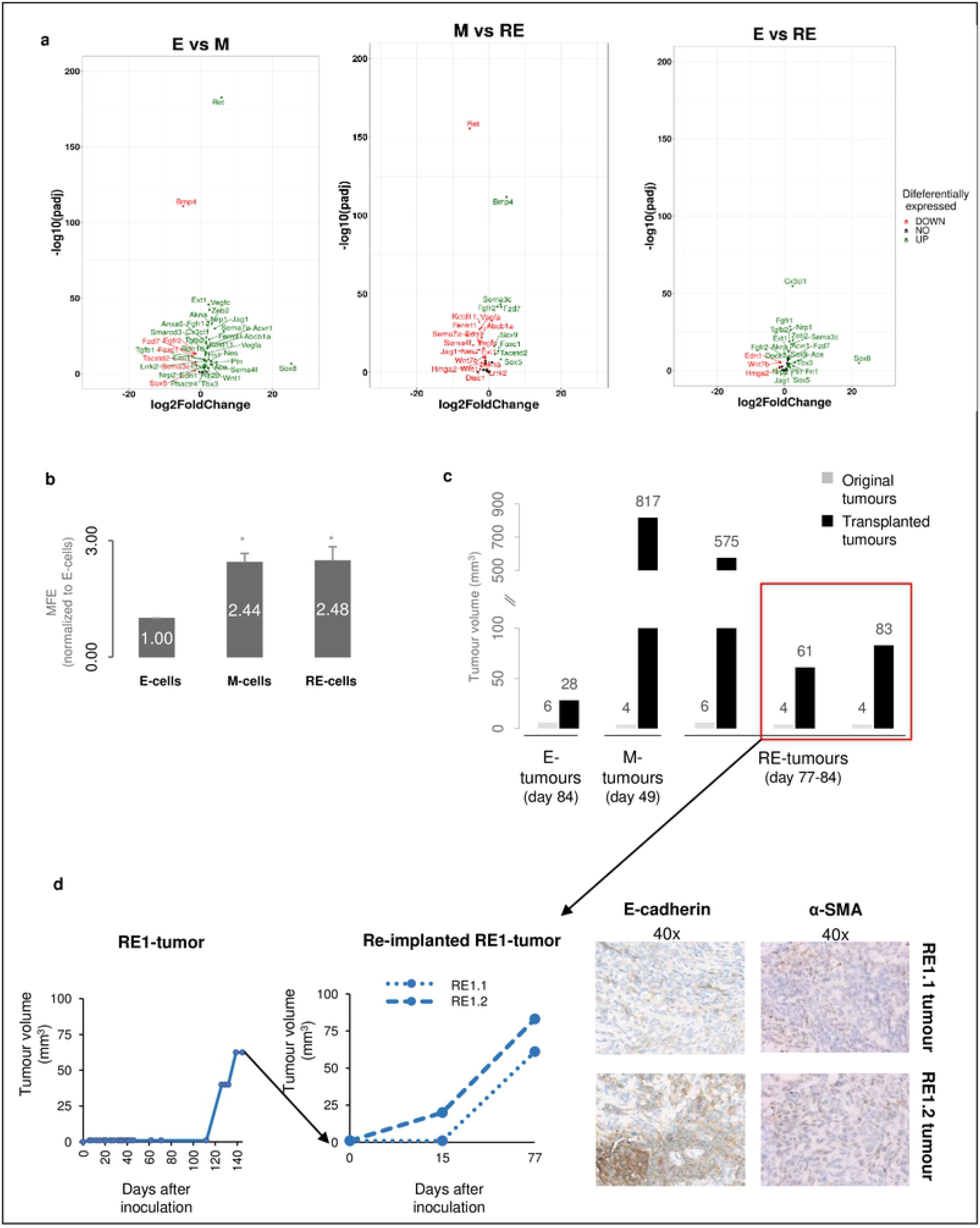
RE-cells display increased stemness potential. **(a)** Volcano plots of stemness-related DEGs. **(b)** E-, M-, RE-cells first-passage mammosphere formation efficiency (* for *p*<5.00E-02). **(c)** Pilot *in vivo* syngeneic transplantation assay for E, M and RE-tumours. Comparison between the final tumour volumes for the original E-, M- and RE-tumours used (grey bars) and the corresponding tumours obtained after syngeneic transplantation (black bars) at the same time-point post-inoculation or transplantation. **(d)** L**eft panel.** Growth curve of a RE-tumour in the first passage in mice (left) and after reinoculation of two tumour pieces (right). **Right panel.** E-cadherin and α-SMA immunohistochemistry staining of representative RE-tumours re-implanted tumours.

Altogether, our results show that RE-cells exhibit a high first-passage mammosphere formation efficiency, which is consistent with the faster growth of tumour transplants *in vivo*.

### RE-cells promote oxidative-phosphorylation after the EMT-related glycolytic shift

To assess whether the similarities in behaviour between M- and RE-cells were associated with equivalent metabolic profiles, we returned to the RNAseq data. Several metabolic pathways were significantly-enriched in the DEG dataset and there were clear differences between E-, M- and RE-cells (Supplementary Fig 5). As a validation, we measured protein expression levels of several key enzymes for glycolysis (HKII [30]), anaerobic respiration (LDH [31]), and oxidative phosphorylation (ND1, NDUFS3 [32]), as well as the rate of lactate production [14] (Fig 6). A significant decreased expression of ND1 and NDUFS3 in M-cells in comparison with E-cells, demonstrated a selective shutdown of the oxidative phosphorylation (OXPHOS) enzymes in EMT (Fig 6d,f). As consequence, pyruvate was diverted towards lactate production, which was demonstrated by an increase in the rate of lactate produced by M-cells (Fig 6e, f). The OXPHOS shutdown observed in M-cells was reverted in RE-cells (Fig 6d-f), however, RE-cells seem to be using lactate-derived pyruvate to feed OXPHOS, unlike E-cells. This can be inferred by HKII downregulation comparing to E-cells (Fig 6b), and increased LDH expression (Fig 6c), also explaining the low ratio of lactate produced in RE-cells (Fig 6e). Altogether, our data show that M-cells displayed a glycolytic metabolism, while RE-cells were strongly committed to revert to OXPHOS, but using its own circuitry that is different from E-cells (Fig 6f). These results shed further light into why RE-cells grow faster than E-cells *in vivo*, since their main source of pyruvate to foster OXPHOS is lactate-derived, which is abundant in the tumoral microenvironment.

**Fig 6.**
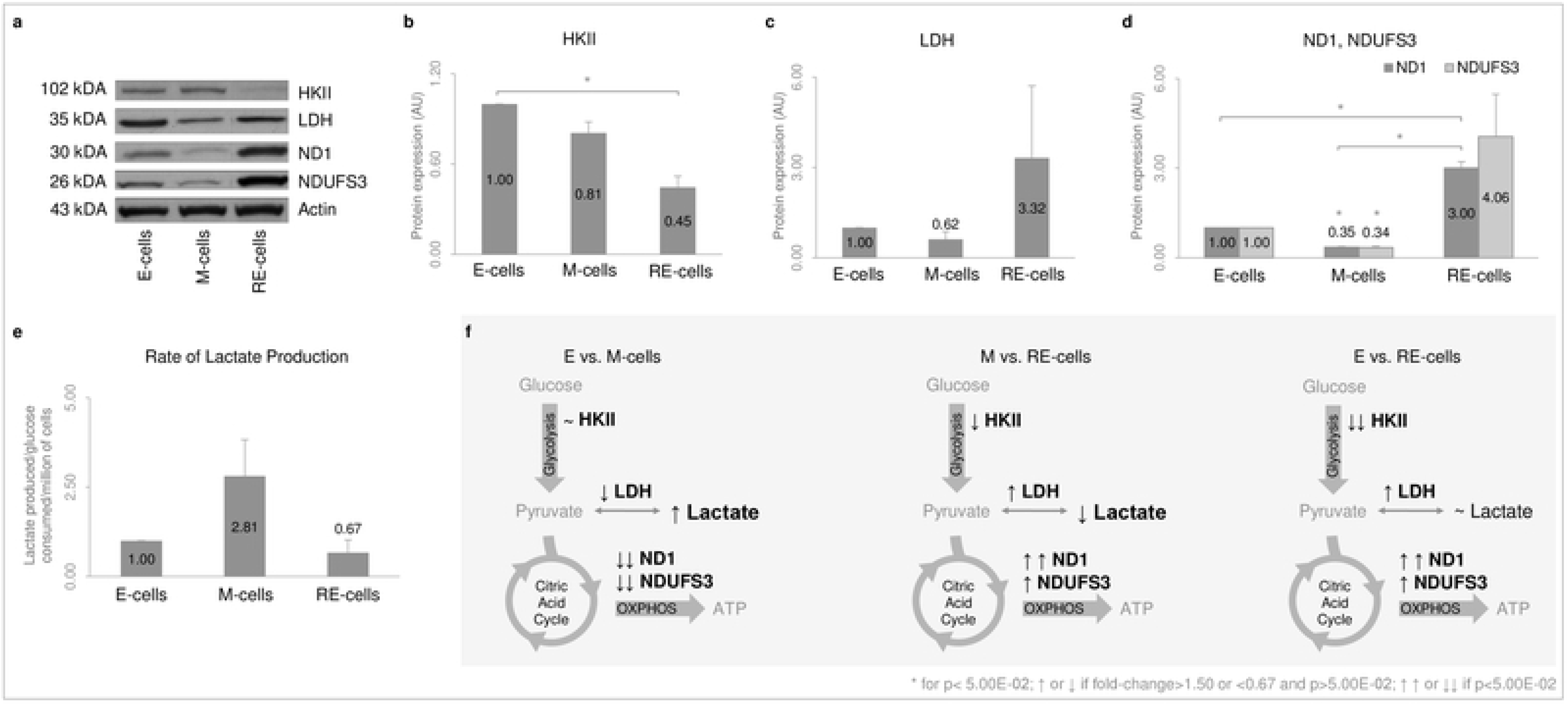
RE-cells display re-activation of oxidative-phosphorylation, after a glycolytic shift observed in M-cells. **(a)** Representative images of western-blot analysis for: HexokinaseII (HKII); Lactate dehydrogenase (LDH); NADH dehydrogenase (ND1); NADH dehydrogenase-ubiquinone-iron-sulfur-protein3 (NDUFS3); Actin (loading-control). **(b)** HKII protein quantification (n=3). **(c)** LDH protein quantification (n=3). **(d)** ND1, NDUFS3 protein quantification (n=3, * for *p*<5.00E-02). **(e)** Rate of lactate produced per glucose consumed per million cells. **(f)** Summary of the expression variation of metabolic enzymes and rate of lactate production for comparisons between E- and M-cells, M- and RE-cells and E- and RE-cells. Each comparison was represented using a scheme portraying aerobic respiration and the variations observed in panels **(b-e)**. Double arrow, *p*<5.00E-02; single arrow, 0.6>fold-change>1.5 and *p*<5.00E-02.

## Discussion

In this study, we show that phenotypic reverted-epithelial cells (RE-cells) do not return to the epithelial state in molecular and functional terms, present mesenchymal features related with aggressiveness and cellular heterogeneity that favour tumour growth *in vivo*. We selected TGFβ1 for EMT induction, as this is a naturally abundant cytokine in tissues, secreted by immune and other cells, which populate the tumour microenvironment [33]. Moreover, TGFβ1 supplementation/withdrawal more closely recapitulates EMT/MET occurrence under physiological conditions, than strategies involving genetic manipulation [34].

We characterized cells that underwent EMT and that presented a phenotypic reversion back to epithelia (RE-cells). RE-cells revealed a distinct transcriptomic profile although resembling E-cells in their cobblestone phenotype and proliferation rate (Supplementary Fig 8). RE-cells displayed mixed E- and M-phenotypic features, were highly heterogeneous *in vitro* with regard to immuno-expression of epithelial and mesenchymal markers, and retained this heterogeneity when growing into large tumours in nude mice. These results are supported by Schmidt et al [35], who suggested that cells undergoing MET may never return to their original epithelial state, gaining aggressive features, and a distinct gene expression profile.

EMT has been associated with increased presence of TICs, tumour progression and aggressiveness [22,36,37], and MET has mainly been associated with increased colonization capacity [8]. So far, neither EMT nor MET have been shown to drive tumourigenesis and the link between EMT plasticity and tumour initiation is still poorly understood. Further, most EMT/MET studies rely on external transformation factors, such as TWIST or SLUG [22,27,38].

Strikingly, we could demonstrate that RE-cells, but not E-cells, could generate large tumours *in vivo*, which grew even larger and faster when transplanted into different animals. These experiments demonstrated that supplementation/withdrawal of a physiologically abundant cytokine was enough to rewire the molecular program of an apparently non-tumourigenic cell line, resulting in increased tumour growth *in vivo* [27]. So far, only one study reported that genetic manipulation of Fra-1 was able to induce EMT and transformation of Eph4 cells, dependent on TGFβ levels [27]. Our work goes beyond that observation, providing evidence that physiological EMT plasticity, without genetic manipulation, may indeed contribute to foster tumour growth.

Unlike the homogenous E or M-cells, RE-cells presented a heterogeneous expression pattern of epithelial and mesenchymal markers (Supplementary Fig 6). Together with our *in vivo* results, these findings recall other EMT-related studies. For example, the work by Tsuji *et al* showed that co-injection of EMT and non-EMT cells originated more aggressive tumours than those obtained by injections of each cell type independently [39]. Upon inoculation, RE-cells were already a mixture of EMT (M-like) and non-EMT (E-like and novel phenotypes) cells, which may, in light of Tsuji *et al*, contribute to the observed RE-tumours increased growth rate. Moreover, the originally homogeneous M-cells also gave rise to high volume M-tumours, similarly to RE-tumours. This suggests that M-cells underwent MET *in vivo*, as M-tumours grew deprived of a persistent TGFβ1 stimulus, unlike M-cells grown *in vitro*. Altogether, and in line with Tsuji *et al*, it is plausible to hypothesize that the intrinsic heterogeneity of RE-cells, and the likely *in vivo* MET in M-tumours, might enable cellular cooperation among distinct subpopulations, promoting tumour growth. Furthermore, RE-cells showed increased self-renewal capacity when compared to E-cells, as shown by increased tumour growth rate upon re-implantation of tumour fragments and by the first-passage mammosphere forming efficiency (Fig 5).

Using this model, we demonstrated that after EMT/MET there is a visible phenotypic reversion back to epithelia, which is not accompanied by molecular and functional reversion, but rather produces a stable heterogeneous cell-population with increased tumorigenic potential. We believe that EMT is crucial to prime E-cells into a more plastic state, culminating, after stimulus removal, into the generation of these distinct subpopulations. Therefore, our model seems to recreate the phenotypic heterogeneity commonly observed in human cancers. A wide range of theories argue that such heterogeneity may arise from clonal evolution, cancer stem-cells, microenvironment cues and/or reversible changes in cancer cells [29,40]. Our work strongly suggests that heterogeneity may be triggered by on/off exposure to a microenvironment cue (i.e. TGFβ1), bestowing cells with aggressive features. In cancer patients, this heterogeneity is likely maintained due to crosstalk between different cancer cell subpopulations, and/or between cancer and non-cancer cell types. Supporting these assumptions is, for example, the direct TGFβ-signalling occurring between platelets and cancer cells, inducing EMT and favouring metastization [41]; the turning on/off of demethylases by different melanoma cell clones, giving rise to a mixture of cells with different tumour growth efficiencies [42]; or the co-existence of epithelial, mesenchymal and hybrid cancer cell states in lung adenocarcinoma providing these tumours with increased survival [37].

In conclusion, our model allowed us to demonstrate that EMT plasticity generates cells with an heterogeneous and unique phenotype, which results in increased stemness and ability to form large tumours *in vivo*, providing evidence that inflammatory cues can influence tumour growth kinetics through EMT/MET transdifferentiation.

## Materials and Methods

### Cell culture

EpH4 [43] provided by Dr. Angela Burleigh and Dr. Calvin Roskelley, cultured as in [25]. MCF10A cell line was cultured in DMEM/F12 Glutamax™ medium, supplemented with horse serum (5%, Lonza), recombinant human insulin (5 ug/mL), penicillin-streptomycin (1%, Invitrogen), hydrocortisone (500 ng/mL, Sigma-Aldrich), cholera toxin (20 ng/mL, Sigma-Aldrich), and recombinant human epidermal growth factor (20 ng/mL, Sigma-Aldrich). Cell authentication by Ipatimup’s Cell Lines Bank, using Powerplex16 STR-amplification (Promega,USA). M-cells obtained as in [25], using Transforming Growth Factor-β1 (TGFβ1, Sigma-Aldrich,USA). RE-cells obtained as in [25] (Supplementary Fig 1).

### RNA extraction/quantification

RNA extraction, cDNA conversion and quantitative-Real-Time-PCR (qRT-PCR) from E-, M-, RE-cells performed as in [25]. qRT-PCR assays used were TaqMan Gene Expression (ThermoFisher,USA) and PrimeTime-qPCR (IDT,USA): CDH1 (Mm00486909_g1), Ocln (Mm.PT.47.16166845), Mgat3 (Mm00483213_m1), Zeb2 (Mm.PT.47.13169136), Vim (Mm01333430_m1), CDH2 (Mm.PT.45.14052292), Twist1 (Mm00492575_m1), GAPDH (Mm99999915_g1), 18S (Hs99999901_s1). Data analysed by 2(-ΔΔCT) method [44] and compared using Mann-Whitney test [45].

### Immunocytochemistry

E-, M-, RE-cells were fixed with methanol (Merck,USA), blocked using 3%BSA-PBS-0,5%Tween20 (Sigma-Aldrich,USA) incubated with anti-Snail (1:50,Cell Signaling,USA), anti-MMP2 (1:50,Calbiochem,USA) and anti-mouse Alexa 488 (1:500,ThermoFisher,USA). Coverslips mounted using Vectashield-DAPI-mounting-medium (Vector Laboratories,USA). Images taken with ZeissImager.Z1AxioCamMRm (Zeiss,Germany).

### Whole Transcriptome Sequencing

RNA for Whole Transcriptome Sequencing (RNAseq) isolated as in [25]. Genomic DNA removed using RNase-Free-DNase (Qiagen,Germany), and purified using RNEasy Mini Kit (Qiagen,Germany). RNA quality analysed using Agilent2100-Bioanalyzer (RIN>9.7). E-, M-, RE-cells sequenced using Illumina Genome Analyzer (n=2) as a service at BCCA. Unique-reads were mapped to NCBI-m37 mouse genome using Bowtie, TopHat2 and differentially-expressed genes (DEGs detected using *edgeR* R package. Genes with log2fold-change>1 or <-1 and corrected p<1.00E-02, were considered DEGs (Supplementary Fig 3). Statistics performed using R. *ClusterProfiler* R package was used for assessment of significantly-enriched GO terms and pathways (*padj*<5.00E-02). The heatmap was performed using the *heatmap.2* function of the *gplots* R package. We used the euclidean method to compute the distance and the complete method to perform the hierarchical clustering.

### E-cadherin and Fibronectin co-immunocytochemistry

E-, M-, RE-cells fixed with methanol (Merck,USA), blocked using 3%BSA-PBS-0,5%Tween20 (Sigma-Aldrich,USA) and co-incubated with anti-E-cadherin (1:50,Cell Signaling,USA), anti-Fibronectin (1:50,Santa Cruz,USA) and anti-rabbit/anti-mouse Alexa 488/594 (1:500,ThermoFisher,USA). Coverslips mounted using Vectashield-DAPI-mounting-medium (Vector Laboratories,USA). Images taken with ZeissImager.Z1AxioCamMRm (Zeiss,Germany).

### BrdU assay

E-, M-, RE-cells were incubated with 1μl of BrdU solution (ThermoFisher,USA) for 90 minutes, washed with Phosphate-Buffered-Saline solution (PBS) and fixed using 4%-formaldehyde (Sigma-Aldrich,USA). Next, cells were treated with 2M-HCl (Merck,USA), washed with PBS-0,5%Tween20-0,05%BSA (Sigma-Aldrich,USA), and incubated with anti-BrdU antibody (1:10, ThermoFisher,USA) and anti-mouse Ig-FITC (1:100,ThermoFisher,USA). Coverslips were mounted using Vectashield-DAPI-mounting-medium (Vector Laboratories,USA). Images were taken with ZeissImager.Z1AxioCamMRm (Zeiss,Germany) and stained-nuclei counted. Statistics performed using Mann-Whitney test [45].

### Focus formation assay

E-, M-, RE-cells plated in 100mm plates and grown for 21 days (TGFβ1-supplemented for M-cells). Brightfield images were taken for phenotype comparison.

### Wound-healing Assay

Wounds produced in confluent E-, M-, RE-cell cultures using a filter-tip. Brightfield images of several E-, M-, RE-cells wounds taken at distinct timepoints (maximum:12h).

### *In vivo* tumourigenesis assay, H&E staining and KI67 immunohistochemistry

1×106/2×106 E-, M- or RE-cells were inoculated in the mammary fat-pad of 5-6 weeks female NIH(S)II-nu/nu mice. Pilot study: 2 mice inoculated with E-, M- or RE-cells in the mammary fat-pad (total n=6). *In vivo* tumourigenicity assay: 5 mice with double E-/RE-cells inoculation and 5 mice with single M-cell inoculation (total n=15). Two mice excluded due to unrelated health issues. *In vivo* syngeneic transplantation assay: 1mm^3^ sections of 1 E-, 1 M- and 3 RE-tumours transplanted into the mammary fat-pad of 5 mice. Tumours measured with callipers and volumes estimated using (Width*Length^2^)/2. Experiments carried out in accordance with European Guidelines for the Care and Use of Laboratory Animals, Directive-2010/63/UE and National Regulation (Diário da República-1.ª série-N.°151). All mice were humanely euthanized. All tumours were formalin-fixed, paraffin-embedded and stained for hematoxylin and eosin. Immunohistochemistry performed for Ki67 as in [46] (ThermoFisher,USA). Three representative fields of each tumour selected and Ki67-stained nuclei counted using D-sight software (A.Menarini Diagnostics, Italy). Sections of 2 E-tumours, 3 M-tumours and 6 RE-tumours conserved in RNA-later (ThermoFisher,USA) used for RNA extraction, cDNA conversion and qRT-PCR as in [25]. Statistics performed using Mann-Whitney test [45].

### First-passage mammosphere formation assay

E-, M-, RE-cells were plated (750 cells/cm2) in 1.2%-polyhema-coated 6-wells (Sigma-Aldrich,USA). Mammosphere growth medium described in [47]. Mammospheres counted after 5 days. Statistics performed using Mann-Whitney test [45].

### Western Blot

E-, M-, RE-cells lysates were immunoblotted as in [46]. The antibodies used were: HexokinaseII (Abcam,UK); Lactate-Dehydrogenase (Santa-Cruz,USA); NADH dehydrogenase1 (Santa-Cruz,USA); NADH-dehydrogenase-ubiquinone-iron-sulfur-protein3 (Abcam,UK); actin (Santa-Cruz,USA). Membranes incubated with horseradish-peroxidase-linked secondary antibodies (GE-Healthcare,UK). Quantification performed using QuantityOne (BioRad,USA). Statistics performed using Mann-Whitney test [45].

### Glucose consumption and lactate production

Glucose GOD-PAD method (Roche,Switzerland) and LO-POD (Spinreact,Spain) measured glucose and lactate in E-, M-, RE-cells conditioned-media. EpH4 culture-medium used for glucose standard curve. Results presented as: lactate produced/glucose consumed/per million of cells. Statistics performed using Mann-Whitney test [45].

## Acknowledgments

We thank Dina Leitão, Regina Pinto, Guilherme Oliveira for assistance with the immunohistochemistry and Ki67-stained nuclei automatic counting. **MS:** Data curation; Formal analysis; Writing – original draft; Writing – review & editing.

## CRedit Author Statement

**MS:** Data curation; Formal analysis; Writing – original draft; Writing – review & editing. **MF:** Data curation; Formal analysis; Investigation; Methodology. **PO:** Conceptualization; Investigation; Methodology; Formal analysis; Writing – original draft. **JC:** Conceptualization; Investigation; Methodology; Formal analysis; Writing – original draft. **SR:** Investigation; Methodology. **MA:** Investigation; Methodology. **AV:** Investigation; Methodology. **DF:** Investigation; Methodology. **AB:** Investigation; Methodology. **CR:** Investigation; Methodology. **JV:** Investigation; Methodology. **JN:** Investigation; Methodology. **AA:** Investigation; Methodology. **AMH:** Investigation; Methodology. **JL:** Conceptualization. **VM:** Conceptualization. **JP:** Conceptualization. **DH:** Conceptualization; Resources. **MFC:** Formal analysis. **CO:** Conceptualization; Resources; Project administration; Supervision; Validation; Funding acquisition; Writing – original draft; Writing – review & editing

